# Reciprocal Modulation Between Amyloid Precursor Protein and Synaptic Membrane Cholesterol Revealed By Live Cell Imaging

**DOI:** 10.1101/328419

**Authors:** Claire E. DelBove, Claire E. Strothman, Roman M. Lazarenko, Hui Huang, Charles R. Sanders, Qi Zhang

## Abstract

The amyloid precursor protein (APP) has been extensively studied because of its association with Alzheimer’s disease (AD). However, APP distribution across different subcellular membrane compartments and its function in neurons remains unclear. We generated an APP fusion protein with a pH-sensitive green fluorescent protein at its ectodomain and a pH-insensitive blue fluorescent protein at its cytosolic domain and used it to measure APP’s distribution, subcellular trafficking and cleavage in live neurons. This reporter, closely resembling endogenous APP, revealed only a limited correlation between synaptic activities and APP trafficking. However, the synaptic surface distribution of APP was inversely correlated to membrane cholesterol levels, a phenomenon that involves APP’s cholesterol-binding site. Mutations within this site not only altered surface APP and cholesterol levels in a dominant negative manner, but also increased synaptic vulnerability to moderate membrane cholesterol reduction. Our results reveal reciprocal modulation of APP and membrane cholesterol levels at synaptic boutons.

## Introduction

Amyloid plaques, one of the pathohistological hallmarks of Alzheimer’s disease (AD), are primarily comprised of β-amyloid peptides (Aβs). Aβs are proteolytic products of the amyloid precursor protein (APP), an integral membrane protein with a single transmembrane domain ^1, 2^. Due to its linkage to AD, APP and its proteolytic processing have been investigated extensively since the early 1990s ^3^. It has been well demonstrated that APP is usually subject to one of two routes of proteolytic processing, amyloidogenic and nonamyloidogenic, catalyzed by three proteases known as the α-, β- and γ-secretases (αS, βS and γS) (**Figure 1A**) ^4^. In the amyloidogenic pathway, APP is first cleaved by βS in its membrane-proximal ectodomain to generate a large soluble protein (sAPPβ) and a membrane-bound C-terminal fragment (CTF) of 99 amino acid residues (C99). Subsequently, C99 is cleaved by γS within the transmembrane domain, yielding Aβs and a short intracellular C-terminal fragment (AICD). In the nonamyloidogenic pathway, APP is first cleaved by αS in the middle of Aβ sequence, yielding a longer soluble ectodomain (sAPPα) and a membrane-bound 83-residue CTF (C83). C83 is also cleaved by γS within the transmembrane domain, generating a shorter 30mer peptide (P3) and an identical AICD ^5^. Interestingly, membrane cholesterol enhances the proteolytic activities of βS ^6, 7^ and γS ^6, 8, 9^ while suppresses αS ^10, 11^. Since APP’s expression is ubiquitous, most studies on APP processing have been conducted in non-neuronal cells ^12, 13^ for technical practicality. In model cell lines, the majority of the APP is found to be located in the intracellular membranes of the Golgi and trans-Golgi network (TGN) and only a small portion is sorted to the plasma membrane ^14^. Those investigations demonstrated that αS cleaves APP in the plasma membrane, whereas both βS and γS cleave APP in endocytic compartments ^15, 16^, suggesting that APP’s subcellular membrane localization determines its proteolytic fate.

**Figure 1.**
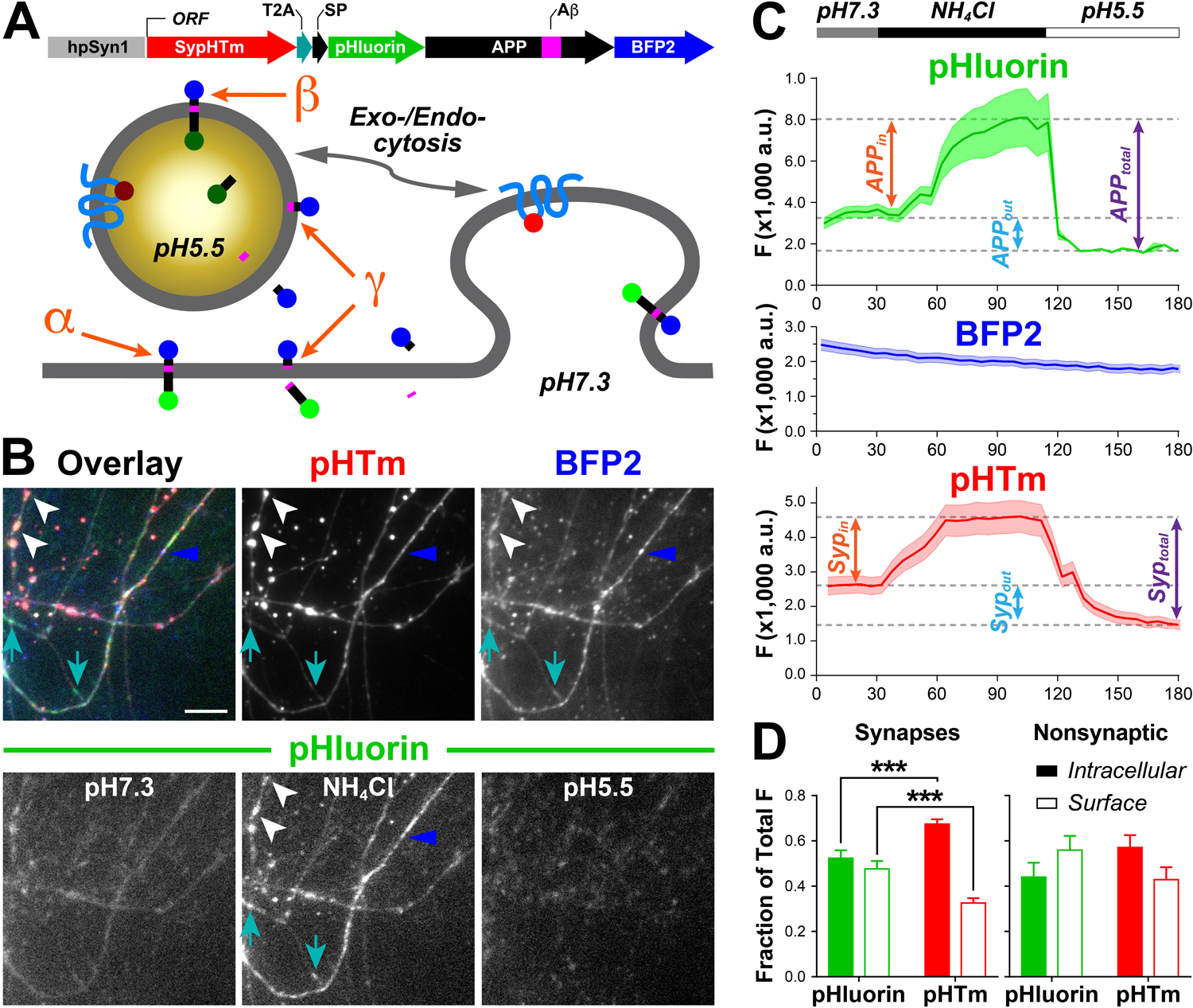
SypHTm:T2A:pH-APP-BFP2 reports APP distribution and marks synaptic vesicles. **A**, the upper diagram illustrates the transgene construct including human Synapsin I promoter (hpSyn1) and an open reading frame (ORF) comprised of Synaptophysin-pHTomato (SypHTm), *thosea asigna* virus 2A peptide (T2A), APP signal peptide (SP), pHluorin (pH), APP containing Aβ, and blue fluorescence protein 2 (BFP2). The lower cartoon demonstrates how extracellular pH and exo-/endocytosis affect pHluorin and pHTm fluorescence and how α-, β-, and γ-secretases cleave pH-APP-BFP2. **B**, top left, overlay of SypHTm (red), pHluorin (green) in 50 mM NH_4_Cl, and BFP2 (blue); top middle, SypHTm in 50 mM NH_4_Cl; top right, averaged BFP2 throughout the course of the experiment; bottom: pHluorin in normal Tyrode’s solution (pH7.3), in 50 mM NH_4_Cl and in pH5.5 Tyrode’s solution. White arrowheads indicate synaptically colocalized SypHTm and pH-APP-BFP2, cyan arrows indicate non-synaptic pH-APP-BFP2, and the blue arrowheads indicate nonsynaptic CTF because of strong BFP2 and weak pHluorin signals. Scale bar, 10 μm. **C**, example of intensity changes of pHluorin, BFP2 and pHTm fluorescence in one FOV (field of view) containing 39 ROIs (regions of interest) during sequential applications of pH7.3 Tyrode’s solution, 50 mM NH4Cl and pH5.5 Tyrode’s solution. Double-ended arrows indicate the calculations of surface, intracellular and total APP and Syp based on fluorescence intensity differences. Shades are s.e.m. **D**, quantification of intracellular (solid bars) and surface (open bars) pHluorin (green) and pHTm (red) fluorescence at synapses (left) and nonsynaptic areas (right). There was a significant difference between pHTm and pHluorin regarding surface or intracellular fractions according to a two-tailed paired t-test (***, *p* = 0.001). No significant difference was found for the nonsynaptic ROIs (two-tailed paired t-test, ns, *p* = 0.0543). Synaptic ROIs, n = 47; nonsynaptic ROIs, n = 23. All error bars represent s.e.m.

However, the use of non-neuronal cells made it difficult to address a key question about APP’s pathological relevance — why it is neurons in the brain that suffer the most in AD? It is worth pointing out that neurodegeneration in the central nervous system (CNS) is the dominant outcome in familial Alzheimer’s disease (FAD) patients bearing APP mutations despite the fact that APP is expressed and cleaved by secretases almost ubiquitously in cells throughout the body. The following reasons further suggest the necessity to study APP in live CNS neurons: **(1)** neurons are morphologically unique for their polarity, extended neurites and intercellular connections known as synapses, which results in an extremely high area and complicated cellular membrane system unmatchable by any other cell types in the body; **(2)** the distinct lipid composition of neuronal membranes (e.g. high cholesterol content) and their unique electrophysiological properties have profound influence on membrane proteins like APP; **(3)** APP is reportedly abundant at synaptic vesicles (SVs), which are clustered at the presynaptic terminals (a.k.a. synaptic boutons) ^17, 18^; **(4)** some hypothetical functions of APP, such as promoting axon outgrowth and synaptogenesis ^19-21^, have yet to be tested in CNS neurons, especially for their pathological relevance; **(5)** APP expression, distribution, and cleavage have been associated with neuronal activity and its proteolytic products are believed to reciprocally affect neurotransmission ^22, 23^; **(6)** APP695, the most disease-relevant APP isoform, is predominantly expressed in neurons but not in astroglia or other types of non-neuronal cells in the CNS ^24^; **(7)** all three major secretases are known to have unique roles in CNS neurons, such as modulation of synaptic transmission and plasticity ^25^; **(8)** Presenilin 1/2 mutations associated with FAD also cause pathological changes similar to those of APP mutations in the CNS ^26^. Interest in neuronal APP has recently surged due to rising debate about the amyloid hypothesis ^27-29^ and also APP’s behaviors unique to neurons^13, 30, 31^.

Because of its spatiotemporal resolution, live cell fluorescence imaging is probably the optimum available approach for tackling the morphological complexity and activity-associated cellular events at submicron-size synapses. Moreover, newly developed fluorescent reporters have enabled the use of fluorescence imaging to qualitatively and quantitatively investigate APP at synaptic boutons. For example, by labeling APP and BACE-1 with two different fluorescent proteins and with two complementary parts of one fluorescent protein, Roy and his colleagues revealed activity-dependent and independent convergence of these two proteins in different intracellular compartments in hippocampal neurons ^30, 31^, behavior that is considerably different from that observed in model cell lines. In another impressive study Groemer *et al* tagged APP’s N-terminal with a pH-sensitive green fluorescence protein (i.e. pHluorin) and was able to quantify the coupling between APP trafficking and SV turnover for the first time ^17^.

Here, we generated pH-APP-BFP2 by adding a pH-insensitive BFP2 at pH-APP’s C-terminal. We also co-expressed a pH-sensitive red fluorescence reporter selective for SVs, Synaptophysin-pHTomato (SypHTm) ^32^, via a bicistronic construct, which enables us to identify synaptic boutons and measure SV turnover. Using these reporters, we performed multichannel live cell imaging to investigate APP’s subcellular distribution, trafficking, association with neuronal activity, and association with membrane cholesterol. Our results not only demonstrated an unexpected correlation between neuronal SV and APP turnover but also revealed reciprocal control between cholesterol and APP, especially in the surface membrane of synaptic boutons.

## Results

### Construction and characterization of a triple-fluorescence reporter system

We started with Groemer’s pH-APP construct (gifted by the J. Klingauf) for several reasons. First, it is based on rat APP695, matching our rat postnatal hippocampal culture. Secondly, its expression is driven by a human Synapsin 1 promoter (hpSynI), ensuring neuron-specific and moderate expression ^33^. Thirdly, pHluorin was inserted behind APP’s short signaling peptide (SP), ensuring the same subcellular distribution pattern as native APP695 ^17, 34^. Because intracellular membrane compartments like SVs and endosomes are often acidic (pH5.5∼6.5) whereas extracellular pH is generally neutral (pH7.35), the pH-sensitive pHluorin located in APP’s ectodomain exhibits fluorescence increase or decrease as APP is externalized or internalized respectively ^17^. However, using pHluorin alone has some major limits. First, it does not allow us to track intracellular APP since pHluorin is quenched. Secondly, N-terminal cleavage by αS or βS detaches pHluorin, yielding non-fluorescent CTFs which are invisible. Thirdly, the only way to quantify total pH-APP is using a high concentration of (e.g. 50 mM) NH_4_Cl to deacidify all intracellular compartments ^17, 35^, which may alter neuronal activity and SV turnover ^36^. We reasoned that adding a pH-insensitive fluorescent protein (i.e. BFP2) to APP’s cytosolic domain (i.e. C-terminal) should address those issues because it provides an independent and constant fluorescent signal for both APP and CTFs. Previous studies have shown that APP fused to fluorescent proteins at both terminals behaves the same as endogenous APP. For example, Villegas et al demonstrated that dual-tagged APPs (i.e. CFP-APP-YFP and FLAG-APP-Myc) were the same as endogenous APP in terms of intracellular distribution and trafficking ^37^. Importantly, neuronal behavior and expression as well as distribution of endogenous APP remain unaltered by the transient expression of those APP fusion proteins ^13, 17, 30, 31, 37^. Hence, we attached BFP2 (gift from Yulong Li) to pH-APP’s C-terminal to generate pH-APP-BFP2. To independently visualize presynaptic terminals and synaptic activity in the same neurons, we inserted Synaptophysin-pHTomato (SypHTm, also a gift from Yulong Li) ^32^ and a viral sequence (T2A) in front of the pH-APP-BFP2 (**Figure 1A**). By so doing, the SypHTm will be co-transcripted with pH-APP-BFP2 in the same mRNA and translated separately with a near 1:1 molar ratio ^38^. Indeed, an immunocytochemistry test using Synaptotagmin I, an SV-specific protein, confirmed that SypHTm was mostly synaptic whereas pH-APP-BFP2 was more evenly distributed across neurites (**Supplementary Figure 1**).

We used a refined protocol ^39^ to achieve ∼30% transfection efficiency in synaptically mature hippocampal cultures (i.e. DIV12-18) ^40^, yielding sparsely labeled neurons, neurites and synapses. To test if those transfected neurons had altered synaptic transmission, we performed whole-cell patch clamp recording on transfected and nontransfected neurons in the same fields of view (FOVs). We observed no differences in the amplitudes and frequencies of spontaneous and miniature excitatory postsynaptic currents (**Supplementary Figure 2A-C** and **D-E** respectively). There was no difference in the paired-pulse ratio either (**Supplementary Figure 2G**&**H**). Hence, we conclude that the expression of our reporters did not affect synaptic transmission. Fluorescence images showed that pHTm was more punctated than BFP2 and pHluorin in the neurites of transfected neurons (**Figure 1B**), consistent with the notion that Syp is an SV-specific protein whereas APP is not. We also observed that most pHTm puncta were overlapped with BFP2 and pHluorin (white arrowheads in **Figure 1B**) but not the other way around (cyan arrows in **Figure 1B**), suggesting the presence of APP and CTFs beyond synapses. There were puncta with strong BFP2 and weak pHluorin fluorescence (blue arrowhead in **Figure 1B**), indicating that either most APP was in acidic compartments or there were more CTFs at those subareas. Next, we asked if the membrane orientation of SypHTm and pH-APP-BFP2 was correct (i.e. pHTm and pHluorin should be located extracellularly and/or luminally) and if we could use them to calculate intracellular and surface fractions of Syp and APP, respectively. To do so, we applied Tyrode’s solution containing 50 mM NH_4_Cl to deacidify all intracellular membrane compartments, followed by pH5.5 Tyrode’s to quench all pHTm or pHluorin (**Supplementary Movie 1**). Applying solutions in this order reduced the concern that neuronal activities were artificially altered ^36^. As expected, we observed significant pHTm and pHluorin fluorescence increases and decreases respectively (**Figure 1B**), which allowed us to measure surface and internal pH-APP-BFP2 or SypHTm (**Figure 1C**). Expectedly, BFP2 fluorescence was stable except for a slow decay likely arising from photobleaching caused by near UV excitation (**Figure 1C**).

First, we calculated **(i)** the total pHluorin and pHTm by subtracting the baseline fluorescence (at pH5.5) from the maximum fluorescence (at 50 mM NH_4_Cl), **(ii)** the surface (out) pHluorin and pHTm by subtracting the baseline fluorescence (at pH5.5) from the pretreatment fluorescence (at pH7.3), and **(iii)** the intracellular (in) pHluorin and pHTm by subtracting the pretreatment fluorescence (at pH7.3) from maximum fluorescence (at 50 mM NH_4_Cl) (**Figure 1C**). Unlike pHluorin, pHTm is only partially quenched at pH5.5, allowing us to identify expressing neurons, neurites and individual synapses without the use of NH_4_Cl ^32, 36^. It was determined that the surface pHluorin fraction was significantly higher than the surface pHTm fraction, while the intracellular pHluorin fraction was significantly lower at synaptic boutons (**Figure 1D**). In comparison, there was little difference between surface and intracellular pHluorin or pHTm (**Figure 1D**), consistent with the notion that Syp is SV-specific whereas APP is a generic membrane protein distributed in both surface and intracellular membranes. The impression that APP was concentrated at presynaptic terminals appears to be due to the presence of hundreds of clustered SVs. Hence, we conclude that SypHTm and pH-APP-BFP2 orient and locate in neuronal membranes in the same way as their endogenous counterparts.

We next asked if pH-APP-BFP2 distributed in the same way as endogenous APP at distal neurites and synapses. We performed fluorescent immunostaining for both endogenous APP and pH-APP-BFP2 using an anti-APP antibody (recognizing APP’s N-terminus) and an anti-GFP antibody (recognizing both pHluorin and BFP2) in nontransfected and transfected cultures respectively. In both cultures MAP2 immunostaining was used to identify dendrites and Synapsin I immunostaining was used to identify synaptic boutons and axons bearing those boutons (**Figure 2A**). The amount of endogenous or exogenous APP in the synapses, axonal shafts, and dendritic shafts was normalized to total APP in the corresponding neurites. We found that endogenous APP and pH-APP-BFP2 were very similar in terms of their distributions in axon, dendrite, and synaptic boutons — more in synaptic boutons, less in dendritic shafts, and least in axon shafts (**Figure 2B**), consistent with the total amount of surface and intracellular membranes in those subcellular structures.

**Figure 2.**
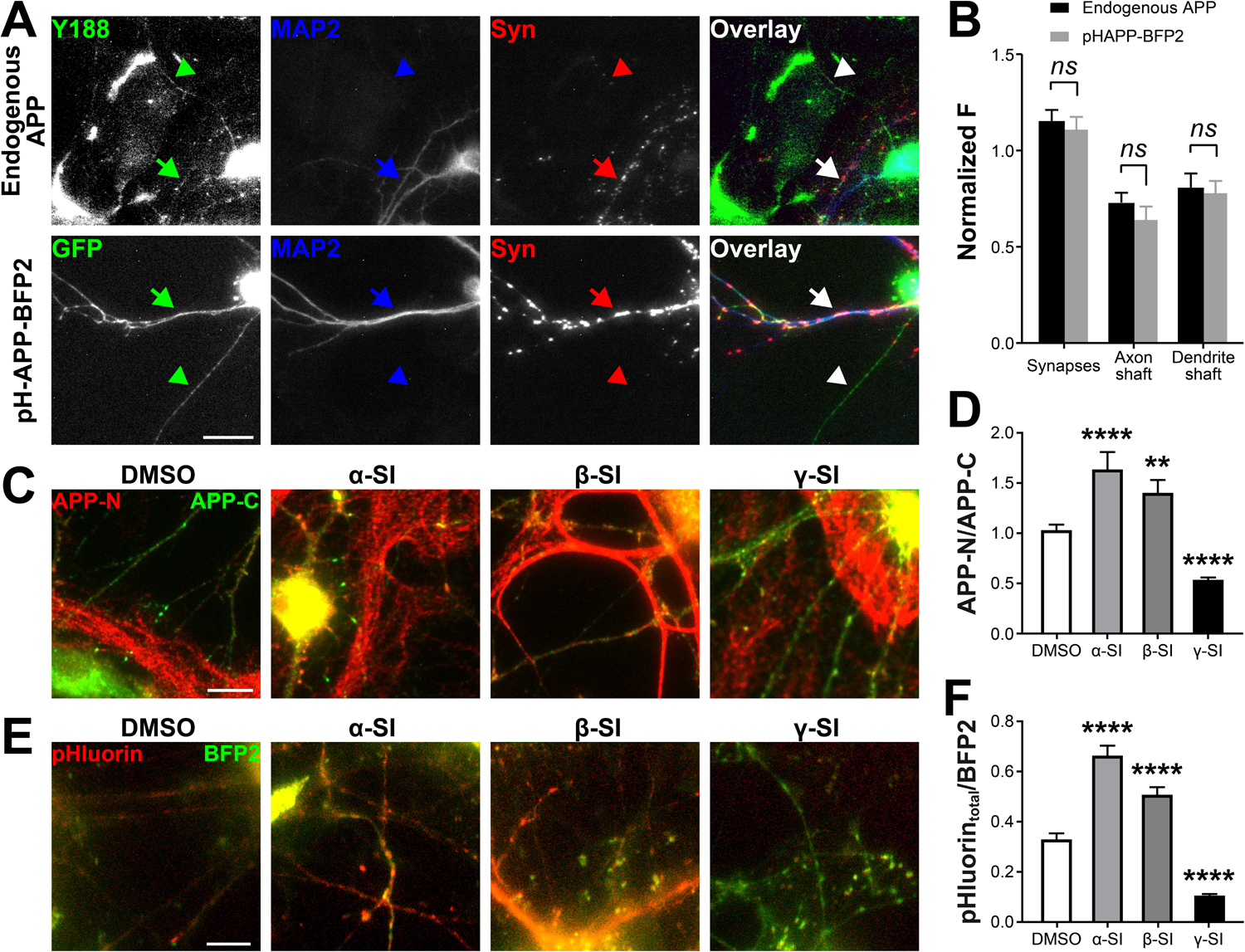
pH-APP-BFP2 is distributed and cleaved in the same way as endogenous APP. **A**, sample images of immunofluorescence labeling of untransfected (top) and transfected cells (bottom). The Y188 antibody was used to detect endogenous APP (green), and anti-GFP antibody was used to detect expressed pH-APP-BFP2 (green). Anti-MAP2 antibody was used to identify dendrites (arrows) and thin neurites with low expression of MAP2 (arrowheads). Anti-Syn was used to identify synaptic boutons. Scale bar, 100 μm. **B,** normalized average APP fluorescence in synaptic boutons, axon shafts and dendritic shafts, normalized to the average fluorescence of the whole processes. Two-way ANOVA detected a significant effect of localization at synapses, axonal shafts and dendritic shafts (F (2, 30) = 25.29, *p* < 0.0001). There was no significant difference between pH-APP-BFP2 and endogenous APP (F (1, 30) = 0.9346, *p* = 0.3414) and no interaction between the two factors (F (2, 30) = 2.256, *p* = 0.1223). **C**, sample images of immunofluorescence labeling for APP N- and C-terminals with and without secretase inhibitors (α/β/γ-SI). Scale bar, 50 μm. **D**, one-way ANOVA detected significant differences in the endogenous APP N/C ratio after α/β/γ-SI treatments (F (3, 380) = 44.153, *p* < 0.0001). Dunnett’s multiple comparisons test showed that α/βSI increased the ratio compared to DMSO (****, *p* = 0.0001; and **, *p* = 0.0014, respectively). γSI significantly decreased the ratio (****, *p* = 0.0001). n_DMSO_ = 160; n_α-SI_ = 21; n_β-SI_ = 38; n_γ-SI_ = 165, where n is the number of Syp-positive puncta (i.e. synaptic boutons). E, sample images of live cell imaging for total pHluorin (for APP’s N-terminal, red color) and total BFP2 (for APP’s C-terminal, green color) after α/β/γ-SI treatments. Scale bar, 10 μm. F, pHluorin/BFP2 ratio after SI treatments. One-way ANOVA detected a significant effect (F (3, 499) = 98.199, *p* < 0.0001) and Dunnett’s multiple comparisons test showed that α-SI and β-SI both increased the ratio compared to DMSO (****, *p* = 0.0001; ****, *p* = 0.0001) while γ-SI decreased it (****, *p* = 0.0001). n_DMSO_ = 132; n_α-SI_ = 126; n_β-SI_ = 80; n_γ-SI_ = 165, where n is the number of ROIs corresponding to Syp-marked synaptic boutons. All error bars represent s.e.m.

Third, we tested if pH-APP-BFP2 was cleaved by the three major secretases in the same manner as endogenous APP was. Based on inhibitor selectivity, potency and usage reported in the literature, we selected GI 254023X (i.e. CAS 260264-93-5), βS inhibitor II (i.e. CAS 263563-09-3) and Compound E (i.e. CAS 209986-17-4) to block αS, βS and γS respectively. Using ELISA assays for sAPPα, we determined that 1 μM 24- hour incubation was effective for GI 254023X to reduce sAPPα levels by more than 50% (**Supplementary Figure 3A&B**). According to Aβ40 ELISA result, 24-hour incubation was also effective for both 0.5 μM βS inhibitor II and 1 μM Compound E, reducing Aβ40 production by more than 30% and 70%, respectively (**Supplementary Figure 3C**). Since αS and βS release APP’s N-terminal ectodomain tagged by pHluorin whereas γS frees the cytosolic C-terminal domain tagged by BFP2, the effect of these secretase inhibitors can be determined by measuring the by N *vs.* C terminal ratio for endogenous APP or the total pHluorin vs. BFP2 ratio for pH-APP-BFP2. Hence, we performed fluorescence immunostaining using antibodies selective for endogenous APP’s N- and C-termini (22C11 and Y188 respectively) and an anti-Syp antibody to mark synaptic boutons (**Figure 2C**). The results showed that αS and βS inhibition increased the synaptic N/C ratio whereas γS inhibition decreased it (**Figure 2D**). For pH-APP-BFP2, we measured the total pHluorin vs. BFP2 ratio at SypHTm-positive synaptic boutons (**Figure 2E**). The observed changes in ratio were nearly identical to those seen for the N/C ratio for endogenous APP (**Figure 2F**). Notably, the fluorescent signals from the live cell imaging were more discrete than those from immunostaining due to its neuron-specific expression, which made the difference among the four groups more significant. Notably, αS inhibitor spared more endogenous APP or pH-APP-BFP2 than βS inhibitor did, consistent with the notion that α-cleavage of APP is more dominant than β-cleavage. Additionally, the changes in the total pHluorin vs. BFP2 ratio at non-synaptic regions were very similar to those at synaptic boutons (**Supplementary Figure 4**). Together, our side-by-side comparisons verified that pH-APP-BFP2 exhibits the same membrane orientation, membrane localization, subcellular distribution and proteolytic processing as endogenous APP.

### APP and synaptic activity

A large body of evidence suggests that intracellular APP translocation and surface turnover are influenced by neuronal activity, linking APP abnormalities to synaptic dysfunction and dementia ^41, 42^. Taking advantage of the fast response of pHluorin fluorescence and the high temporal resolution of optical imaging, we asked if APP exhibited synaptic activity-associated changes in subcellular distribution, a question that cannot be readily addressed using conventional biochemical approaches in nonneuronal cells or fixed tissues. First, we examined if the intra-neurite translocation of APP and/or CTFs was affected by synaptic activity. We applied two discrete stimuli (i.e. a 1-minute 10-Hz electric field stimulation and a 1-minute 90 mM K+ perfusion) to simulate high-frequency firing and prolonged depolarization. The two stimuli were separated by a 1-minute resting period long enough for the synapses to recover. We monitored the BFP2 signal instead of the pHluorin signal in order to track both surface and intracellular APP, as well as CTF (**Figure 3A** and **Supplementary Movie 2**). Again, SypHTm was used to mark synaptic boutons along neurites. The selected BFP2 kymograph exemplifies the diverse trafficking behavior of APP and CTF, i.e. stationary or mobile, anterograde or retrograde, and toward or away from nearby synaptic boutons (**Figure 3B**). In comparison, SypHTm kymograph showed that synaptic boutons were stationary. When plotted against time, neither the velocity (**Figure 3C**) nor the distance from the nearest synapse (**Figure 3D**) of the moving BFP2 puncta exhibited any significant difference during stimulation versus resting periods, suggesting that APP and/or CTFs move along distal neurites rather randomly and are not influenced by acute changes in synaptic activity.

**Figure 3.**
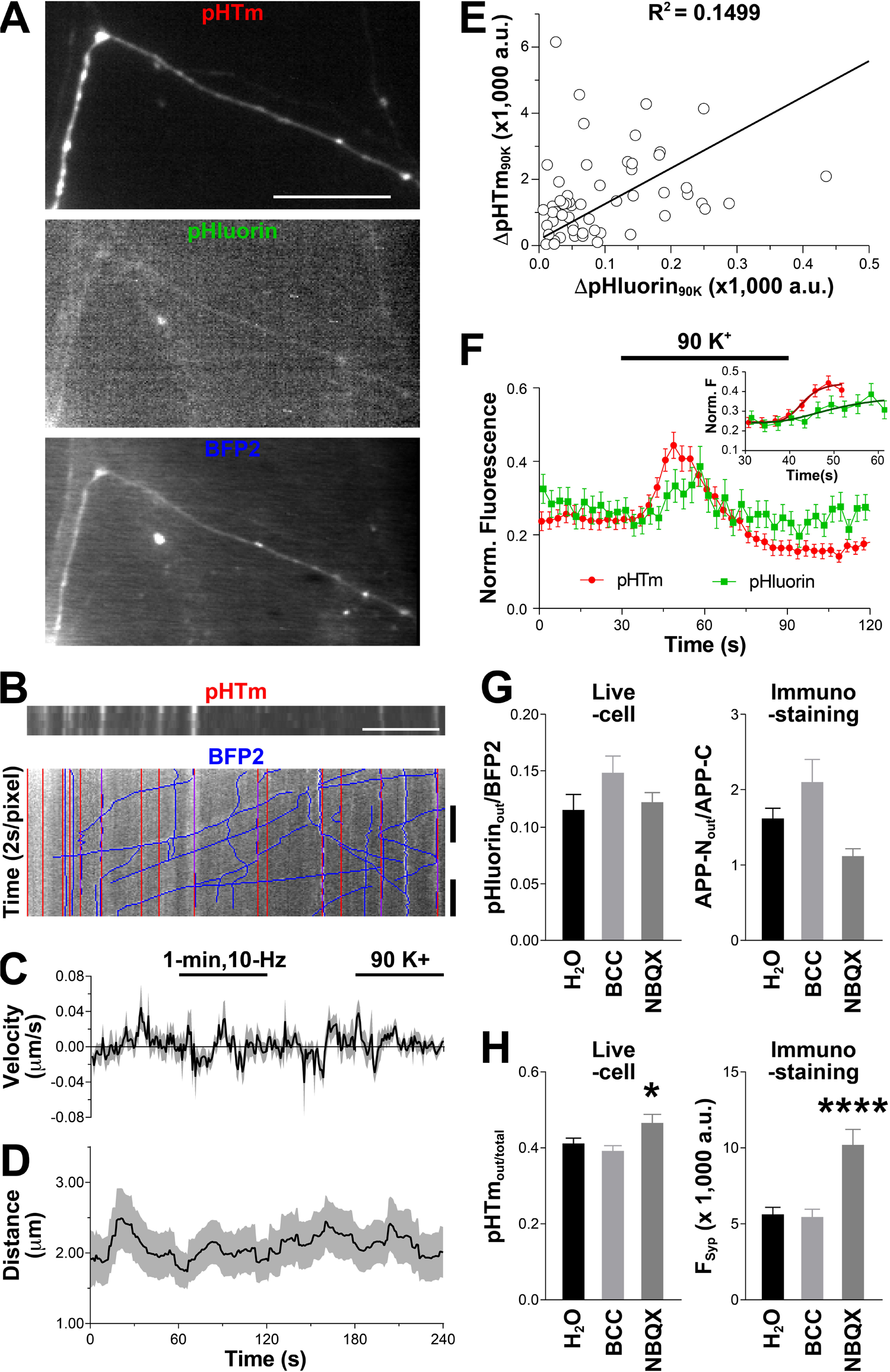
APP trafficking is not activity-associated. **A**, sample images of a distal neurite. Scale bar, 10 μm. **B**, sample kymographs that show the changes in fluorescence over time during 10 Hz field stimulation and 90K stimulation along the process shown in **A**. BFP2 was imaged at 0.5 Hz, while pHTm was imaged every minute. Hand drawn BFP2 tracks are shown in blue, stationary synaptic boutons are in red, and purple represents overlap. Scale bar, 10 μm. **C**, mean velocity of the moving BFP2 puncta. Movement towards the nearest synapse was defined as positive. **D**, mean distance from the moving BFP2 puncta to the nearest synapse. For **C-D**, only visible puncta contribute to the mean and s.e.m, with at most 81 puncta and at least 50 puncta at any given time. Totally, 241 tracks are taken in 13 kymographs from 4 trials. **E**, ΔpHTm_90K_ and ΔpHluorin_90K_ represent the maximal increase of fluorescence of every ROI during 90 mM K^+^ perfusion. n = 56 ROIs from 3 trials. The line shows a linear regression of slope 1.25 ± 0.36. Pearson’s R^2^ = 0.1499. **F**, synaptic pHluorin and pHTm fluorescence normalized to the maximal values set by 50 mM NH_4_CL. The pHluorin increase during stimulation with 90 mM K^+^ is significantly smaller than that of pHTm according to an unpaired two-tailed *t*-test (p = 0.0224). n = 56 ROIs from 3 FOV. The inset shows variable slope (4 parameters) curves fits to the rising phases of pHTm and pHluorin (30 - 60s) with the constraints Top = 0.4437 or Top = 0.3864, which are the maximums of pHTm and pHluorin, respectively. Based on the fittings, for pHTm, t_1/2_ = 12.98 s; for pHluorin, t_1/2_ = 20.15 s. **G**, one-way ANOVA detected significant difference in surface pHluorin vs. BFP2 ratio for pH-APP-BFP2 (left) (F (2, 352) = 3.6597, *p* = 0.0267). However, Dunnett’s multiple comparisons test showed no significant difference between H_2_O and bicuculine (BCC) and between H_2_O and 2,3-dihydroxy-6-nitro-7-sulfamoyl-benzoquinoxaline (NBQX), (p = 0.0741 and 0.9941, respectively). n_H2O_ = 80; n_BCC_, n = 110; n_NBQx_ = 165 synaptic boutons. One-way ANOVA detected significant difference in surface APP N-terminal vs. total APP C-terminal immunostaining for endogenous APP (right) (F (2, 447) = 8.2663, *p* = 0.0003). However, Dunnett’s multiple comparisons test showed no significant difference between H_2_O and BCC and NBQX, (p = 0.5406 and 0.0117, respectively). All three n = 150 synaptic boutons selected randomly. **H**, one-way ANOVA detected significant difference in surface vs. total pHTm ratio (left) (F (2, 352) = 4.3003, *p* = 0.0143). Dunnett’s multiple comparisons test showed no significant difference between H_2_O and BCC and significant difference between H_2_O and NBQX, (p = 0.3157 and 0.0084, respectively). Same n as (**G**). One-way ANOVA detected significant difference in immunostaining for endogenous APP (right) (F (2, 447) = 19.9310, *p* < 0.0001). Dunnett’s multiple comparisons test showed no significant difference between H_2_O and BCC and significant difference between H_2_O and NBQX, (p = 0.8307 and *p* < 0.0001, respectively). Same n as (**G**). All shadows (**C, D**) or error bars (**F, G** & **H**) represent s.e.m.

Secondly, we asked if surface-internal turnover of APP is correlated to activity-associated SV exo-/endocytosis at synaptic boutons, which can be monitored via the SypHTm signal ^32^. If APP is enriched in releasable SVs, SypHTm and pHluorin fluorescence should exhibit coordinated changes during stimulation. We applied 90 mM K^+^, and simultaneously measured the pHTm and pHluorin fluorescence changes. While we did observe an increase of pHluorin fluorescence at synaptic boutons, the amount of increase was not proportional to that of pHTm fluorescence (**Figure 3E**), suggesting that the APP externalization and SV release were discordant. Next, we analyzed the time courses of fluorescence fluctuations for pHluorin and pHTm during and after the stimulation. Again, there was a clear discrepancy between the two, namely pHluorin fluorescence changes lagged behind that of pHTm (**Figure 3F**), further suggesting that most APP was not located in readily releasable SVs and thus was not recycled along with SVs. Moreover, we did not observe strong correlations for the total, surface, or intracellular pHluorin and pHTm fluorescence intensities (**Supplementary Figure 5**), again disputing APP and Syp colocalization at the same surface or intracellular membrane at presynaptic terminals. Together, it is safe to say that unlike Syp, APP is not an SV protein.

We further probed if prolonged network activity change alters the synaptic localization and surface-internal trafficking of APP. To do so, we globally enhanced or suppressed neuronal network activity by blocking GABAergic (inhibitory) or glutamatergic (excitatory) neurotransmission with 10 μM bicuculline (BCC) or 10 μM 2,3-dihydroxy-6-nitro-7-sulfamoyl-benzoquinoxaline (NBQX) for two hours ^43^. Immediately after the treatments, cells were either subjected to live cell imaging or fixed for immunostaining. In comparison to a sham control (i.e. H_2_O), neither BCC nor NBQX treatment changed the surface APP fraction based on the measures of both pH-APP-BFP2 or endogenous APP (**Figure 3G**). Moreover, ELISA measurements detected no difference in the production of Aβ40, Aβ42 or sAPPα(**Supplementary Figure 6**), further ruling out activity-associated changes of APP cleavage. In contrast, the surface vs. total SypHTm was significantly increased after NBQX treatment (**Figure 3H**), which was expected due to presynaptic scale-up ^43, 44^. Together, these data demonstrated that, other than a limited and delayed correlation between acute synaptic activity and the surface-internal trafficking of APP, neuronal activity has little impact on APP trafficking and turnover.

### Synaptic surface membrane cholesterol and APP distribution

APP’s correlation to cell membrane density prompted us to ask if APP is related to major membrane components important for presynaptic terminals. Cholesterol is one of the most abundant and essential lipids in the presynaptic terminals ^18, 45^. Recently, structural studies using nuclear magnetic resonance spectroscopy (NMR) demonstrated that APP interacts with cholesterol through a partly membrane-buried cholesterol-binding site ^46-48^. Moreover, cholesterol has previously been linked to AD in various ways. First, ApoE4, the highest genetic risk factor for late-onset AD (a.k.a. sporadic AD), leads to a reduced cholesterol supply to neurons compared to the AD-protective or benign ApoE2&3 variants ^49^. Second, APP and ApoE reciprocally modulate each other’s expression ^50-52^. Third, when cholesterol transportation to the plasma membrane is disrupted by genetic defects like NPC1 (found in Niemann-Pick Type C1 disease), AD-like histopathologies, including abnormal Aβ metabolism, neurofibrillary tangles and neurodegeneration, appear ^53^. Fourth, increasing membrane cholesterol shifts APP processing from the non-amyloidogenic mode to amyloidogenic by suppressing αS ^10, 11, 54^ and promoting βS and γS ^55, 56^. Fifth, age-related loss of membrane cholesterol is often accompanied by synaptic dysfunction during the preclinical stage of AD ^57^. Sixth, a deficiency in synaptic membrane cholesterol impairs synaptic plasticity, the biological basis of learning and memory ^58^. Therefore, we decided to examine the association between membrane cholesterol and APP.

We asked if and how a moderate reduction of membrane cholesterol would affect synaptic APP processing and trafficking. We used an empirically-determined 90-minute 1 mM MβCD (methyl-β-cyclodextrin) treatment to reduce cholesterol levels in the synaptic membranes (**Supplementary Figure 7**A). Based on Filipin staining of membrane cholesterol, this treatment caused a ∼19% reduction of absolute Filipin fluorescence or ∼10% after adjusting for membrane density (please see method section for details) (**Supplementary Figure 7B-D**). Importantly, no detectable membrane damage or morphological change occurred at distal neurites or synaptic boutons (**Supplementary Figure 7B**). Using live cell imaging, we observed that this mild MβCD treatment caused a significant decrease of total pHluorin fluorescence but only an insignificant decrease of BFP2 (**Figure 4A-C**), leading to a significant reduction in the ratio of total pHluorin to BFP2 (**Figure 4D**). These results indicate that the mild MβCD treatment led to enhanced α-cleavage of APP in the surface membrane but had little change on γ-cleavage. Indeed, the relative fraction of surface APP was significantly increased (**Figure 4E**), which could be due either to enhanced cleavage of intracellular APP or elevated transportation of intracellular APP to cell surface. Similar changes in total and surface pHTm signals were observed, although they were less significant (**Figure 4F&G**).

**Figure 4.**
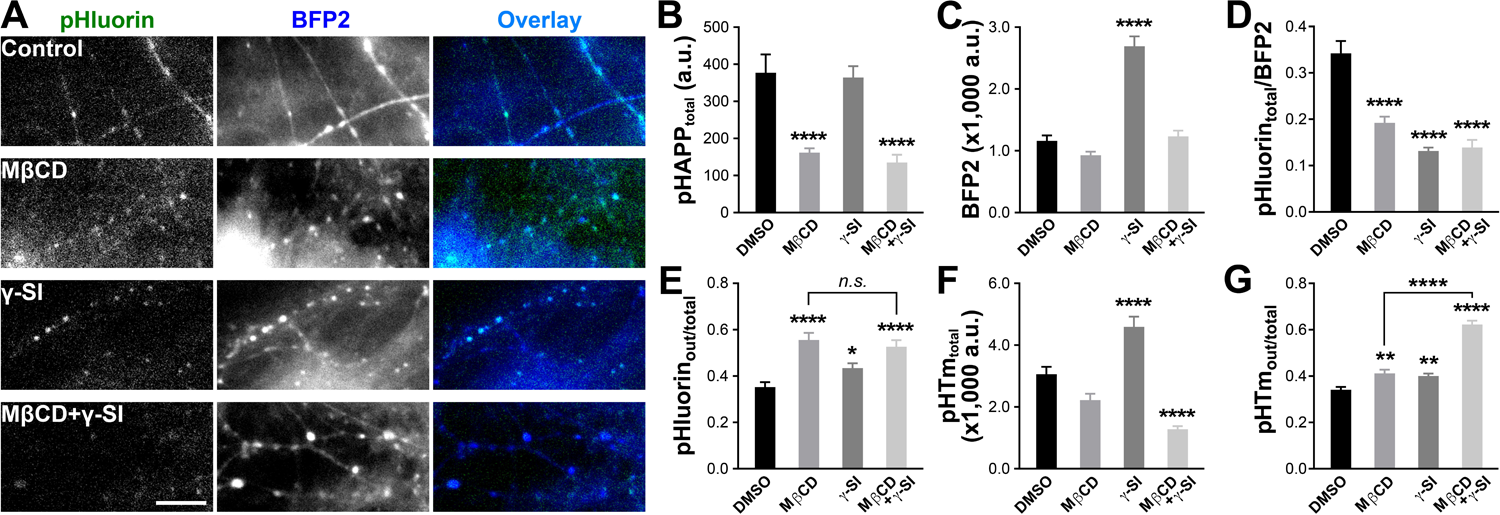
Membrane cholesterol affects the distribution and cleavage of APP. **A**, sample images of pHluorin, BFP2 and their overlays with four different treatments. Scale bar, 10 μm. **B**, average total pHluorin fluorescence intensities in synaptic boutons marked by SypHTm. One-way ANOVA detected significant differences among treatments (F (3, 360) = 14.93, *p* < 0.0001). Dunnett’s multiple comparisons test showed that MβCD significantly decreased total pHluorin fluorescence without or with γ-SI (****, *p* = 0.0001 for both). However, γ-SI alone made no difference (p = 0.9834). **C**, average BFP2 fluorescence intensities in synaptic boutons marked by SypHTm. Oneway ANOVA detected significant differences among treatments (F (3, 360) = 48.65, *p* < 0.0001). Dunnett’s multiple comparisons test showed that only γ-SI significantly increased BFP2 fluorescence compared to WT (****, *p* = 0.0001). **D**, average total pHluorin vs. BFP2 ratio in synaptic boutons marked by SypHTm. One-way ANOVA detected significant differences among treatments (F (3, 360) = 35.04, *p* < 0.0001). Dunnett’s multiple comparisons test showed that MβCD, γ-SI and MβCD+γSI significantly decreased the ratio (****, *p* = 0.0001). **E**, average surface vs. total pHluorin ratio in synaptic boutons marked by SypHTm. One-way ANOVA detected significant differences among treatments (F (3, 360) = 12.9832, *p* < 0.0001). Dunnett’s multiple comparisons test showed that both MβCD and γ-SI significantly increased the ratio (****, *p* = 0.0001; *, *p* = 0.0343, respectively), as did the combination (****, *p* = 0.0001), but there is no additive effect for MβCD and γ-SI combined compared to MβCD alone (two-tailed unpaired *t*-test, *p* = 0.4855). **F**, average total pHTm in synaptic boutons marked by SypHTm. One-way ANOVA detected significant differences among treatments (F (3, 360) = 30.86, *p* < 0.0001). Dunnett’s multiple comparisons test showed that γ-SI significantly increased total pHTm (****, *p* = 0.0001), whereas MβCD and MβCD+γ-SI decreased it (n.s., *p* = 0.0888; ****, *p* = 0.0001, respectively). **G**, average surface vs. total pHTm in synaptic boutons marked by SypHTm. One-way ANOVA detected significant differences among treatments (F (3, 360) = 76.19, *p* < 0.0001). Dunnett’s multiple comparisons test showed that MβCD, γ-SI and MβCD+γ-SI significantly increased the ratio (**, *p* = 0.0014; **, *p* = 0.003; ****, *p* = 0.0001, respectively). Additionally, a two-tailed *t*-test showed a significant difference between MβCD+γ-SI and MβCD (****, *p* < 0.0001). For **B-G**, n_DMSO_ = 90 (FOV = 5); n_MβCD_ = 75 (FOV = 5); n_γ-SI_ = 119 (FOV = 7); n_MβCD+γ-SI_ = 80 ROIs (FOV = 4), where n is the number of ROIs corresponding to SypHTm-marked synaptic boutons.

Since cholesterol modulates different secretases differently and since different secretases cleave APP at different membrane compartments, is it possible that the increase in surface APP fraction was an indirect effect due to cholesterol-dependent change of secretase activity? For the reasons mentioned above, we can exclude αS. βS is also unlikely because it cleaves intracellular APP and is suppressed by MβCD. Inhibition of γS did increase surface pHluorin and SypHTm fractions in addition to the accumulation of CTF (i.e. increase of BFP2 fluorescence) (**Figure 4B-G**). However, the lack of change in BFP2 fluorescence (**Figure 4C**) suggests that our mild MβCD treatment was insufficient to suppress the γ-cleavage of APP. Since γS can also be present in the plasma membrane and can form a complex with αS ^12^, we considered whether it is possible that our MβCD treatment suppressed αS via attached γS. To test this, we applied a γS inhibitor. Again, inhibition of γS significantly increased BFP2 fluorescence without altering total pHluorin, leading to a decreased total pHluorin vs. BFP2 ratio (**Figure 2F** and **4B-D**). However, the surface APP increase by γS inhibition, although significant in comparison to DMSO control, is much less than that of MβCD (**Figure 4E**), indicating different underlying mechanisms. We then tested the combination of both MβCD and application of a γS inhibitor. The fluorescence changes of pH-APP-BFP2 were very similar to those caused by MβCD alone (**Figure 4B-E**), whereas the changes of SypHTm were different from either γS inhibition or MβCD alone (**Figure 4F&G**). These results suggested that MβCD and γS inhibition impact the surface APP distribution through different mechanisms and that the effect of MβCD treatment was predominant. It is likely that MβCD and γS act on SVs turnover through different mechanisms as well.

### APP’s cholesterol-binding affects its surface distribution

If membrane cholesterol modulates APP surface distribution independent of secretases, does that involve a direct interaction between APP and cholesterol in cell membranes? Recently, a cholesterol-binding motif overlapping with APP’s transmembrane domain has been identified using NMR ^46-48, 59^. The physiological relevance of this binding site has yet to be determined. We therefore introduced two different point mutations (G700A and I703A) into pH-APP-BFP2 separately. Both of them are outside of the major cleavage sites of α/β/γ-S and strongly reduce APP’s affinity for cholesterol ^46^. We expressed both wildtype (WT) and mutant forms of SypHTm:T2A:pH-APP-BFP2 in cultured neurons (**Figure 5A**). While the expression levels varied, the proteolytic cleavages of all three forms were very similar according to the ratio of total pHluorin *vs*. BFP2 (**Figure 5B**), confirming that both the cholesterol binding site mutations did not affect secretase cleavage. Intriguingly, both mutants caused a significant increase in the surface fraction of APP (**Figure 5C**), suggesting that cholesterol-binding to APP is involved in restricting neuronal surface APP distribution. At the same time, there was an insignificant increase of surface SypHTm fraction (**Figure 5D**), again disputing a strong link between SVs and APP.

**Figure 5.**
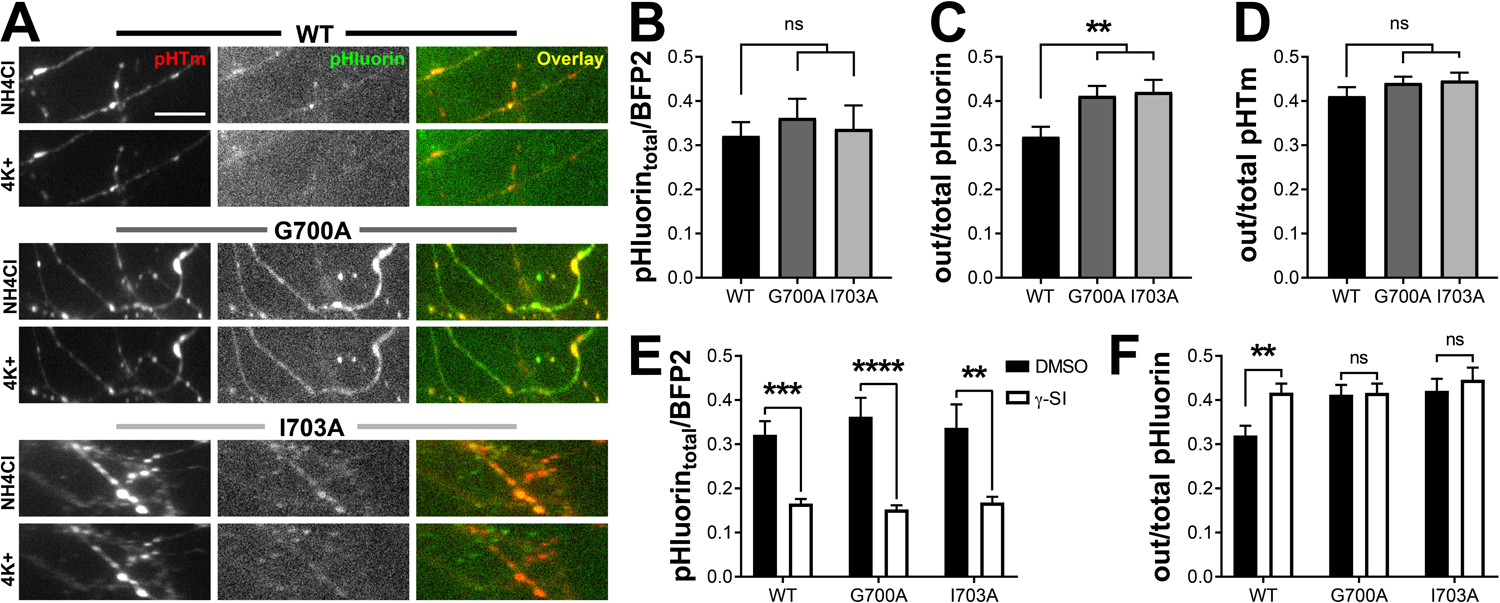
Point mutations within APP’s cholesterol-binding motif increase APP surface fraction. **A**, sample images of pHTm, pHluorin and their overlays under 50 mM NH_4_Cl or 4K+ normal Tyrode’s solution. Scale bar, 10 μm. **B**, neither mutation affects the ratio of total pHluorin to BFP2 at the synapses (one-way ANOVA, F (2, 245) = 0.2601, *p* = 0.7712, Dunnett’s multiples comparisons test: WT vs. G700A, ns, *p* = 0.6976; WT vs. I703A, ns, *p* = 0.9539). **C**, G700A and I703A mutations cause an increase in the surface fraction of pHluorin at the synapses (one-way ANOVA, F (2, 245) = 5.652, *p* = 0.0040, Dunnett’s: WT vs. G700A *p* = 0.008; WT vs. I703A *p* = 0.0077). **D**, the pHTm surface fraction is unchanged by the introduction of mutations to pH-APP-BFP2 (one-way ANOVA, F (2, 245) = 1.267, *p* = 0.2835). **E**, both mutants are affected by the γ-SI, demonstrating that γ-secretase is able to cleave them. Two-way ANOVA indicated that only application of the secretase inhibitor affected the pHluorin total to BFP2 ratio. In summary, there was no effect due to mutation (F (2, 531) = 0.1346, *p* = 0.8741), significant variance from γ-SI treatment (F (1, 531) = 54.6, *p* < 0.0001), and no interaction (F (2, 531) = 0.5277, *p* = 0.5903). The results of Sidak’s multiple comparisons test comparing only the untreated and the γ-SI treated samples within each APP variant are shown on the plot (WT, ***, *p* = 0.0002; G700A, ****, *p* < 0.0001; I703A, **, *p* = 0.0013). **F**, Two-way ANOVA was used to investigate the effects of γ-inhibition on the mutants. Although the interaction between APP sequence and γ-SI treatment did not quite reach significance with α = 0.05 (interaction F (2, 531) = 2.421, *p* = 0.0898), when Sidak’s multiple comparisons test was used only to compare vehicle control to γ-SI, γ-SI significantly increased the surface fraction of the wild-type (p = 0.0062) but not the mutants (G700A, *p* = 0.9984 and I703A, *p* = 0.8847). ANOVA confirmed that γ-SI treatment (F (1, 531) = 4.791, *p* = 0.0290) and cholesterol-binding deficiency (F (2, 531) = 4.022, *p* = 0.0185) cause significant alterations in pHluorin surface fraction unlikely to occur by chance when the entire data set is considered. For B-F, n is the total number of synapse ROIs from at least 3 experiments for every condition, and the data set is the same: n_WT_ = 84, n_G700A_ = 96, n_I703A_ = 68, n_WT+γ-SI_ = 112, n_G700A+γ-SI_ = 114 and n_I703A+γ-SI_ = 63. All error bars represent s.e.m.

Next, we tested if the altered surface distributions of those mutants were the results of changes in γ-cleavage, which could be affected by APP’s affinity to cholesterol. Again, we used γS inhibitor and observed increased BFP2 fluorescence without significant change in total pHluorin fluorescence (**Supplementary Figure 8**), which resulted in a significant decrease of total pHluorin vs. BFP2 ratio in both WT and mutants (**Figure 5E**). Hence, it is unlikely that either point mutation interferes with γ-cleavage. Notably, while γS inhibition significantly increased the surface APP fraction in WT as we previously observed, it had no effect on either the G700A or I703A mutant (**Figure 5F**). There could be two possibilities. One was that those mutations modified the surface APP distribution via a cholesterol-dependent but γS-independent mechanism, and the other was that these mutations simply masked the effect of γS inhibition. The latter is deemed unlikely as both mutations were outside of the range of γ-cleavage sites and exhibited unaltered γS cleavability. We therefore conclude that the binding of cholesterol to APP either prevents the protein from being transported to surface membrane or promotes its internalization independent of SV turnover at presynaptic terminals.

### Binding of cholesterol by APP is important for presynaptic integrity

Since APP regulates membrane cholesterol and since its cholesterol-binding motif is involved, we asked whether and how the two APP mutants will affect cholesterol concentration in neuronal membranes including synaptic boutons. First, we transfected cells with WT or mutation-bearing SypHTm:T2A:pH-APP (C-terminal BFP2 was removed because its emission spectrum overlaps with Filipin’s). We also preloaded the cells with AM1-43, a fixable variant of FM1-43, which we used to normalize the Filipin signal for the variation of membrane density. We performed Filipin staining at 4°C to reduce permeabilization and limit the staining to the neuronal surface membrane as much as possible (**Figure 6A**). During analysis, we used the pHTm signal to identify transfected neurons and their processes, which were divided into regions of interest that also encompass synaptic boutons. Intriguingly, there was significantly less membrane cholesterol in the neurites and synaptic boutons of I703A-expressing neurons based on absolute as well as normalized Filipin fluorescence intensity (**Figure 6B1**). The normalized Filipin results also suggested significantly lower membrane cholesterol concentration in the neurites and synaptic boutons of G700A-expressing neurons (**Figure 6B2**). Notably, the effect of those two mutants on membrane cholesterol was dominant-negative since endogenous WT APP was still present in those transfected neurons.

**Figure 6.**
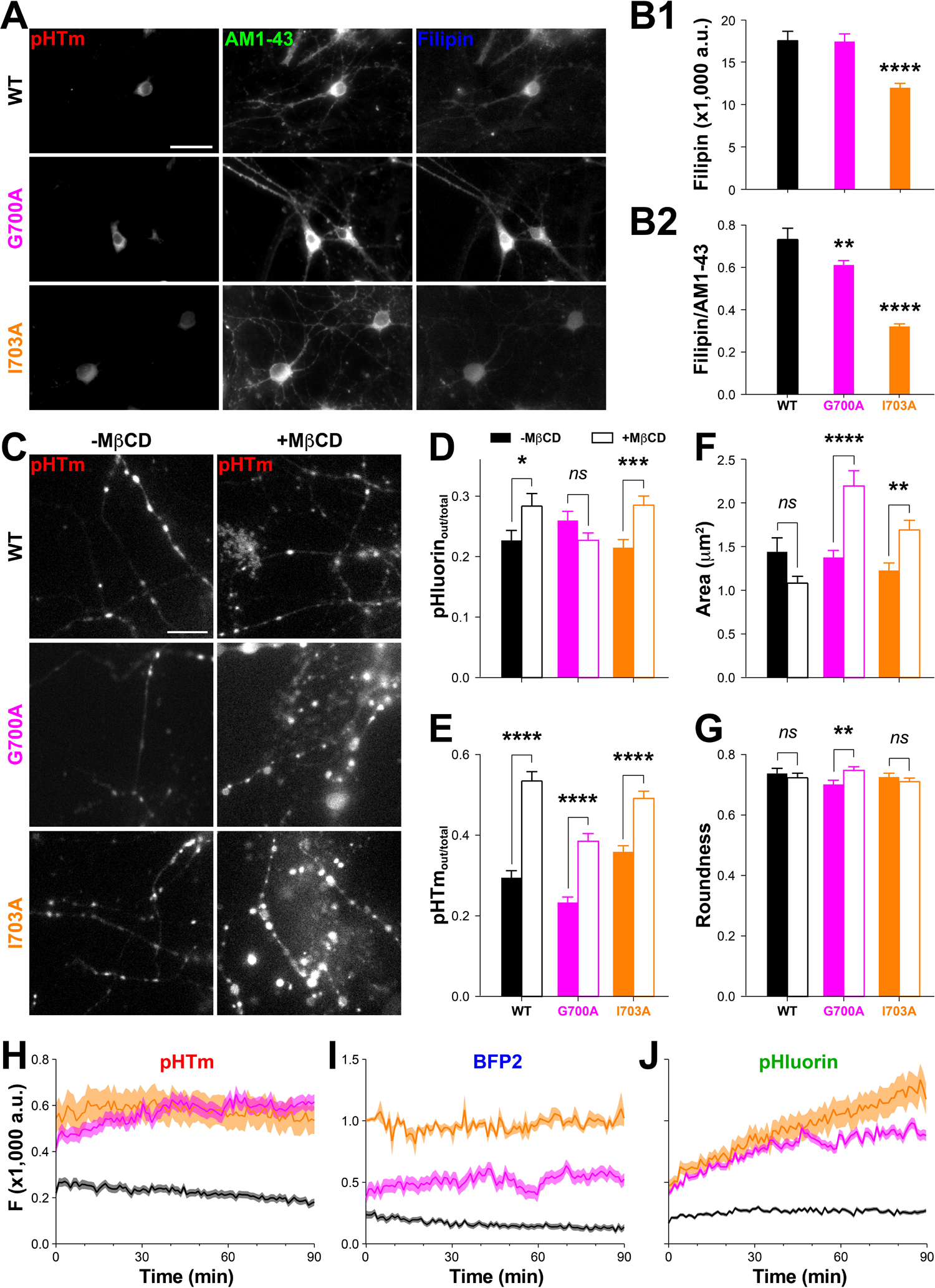
Point mutations in the cholesterol-binding motif render presynaptic terminals vulnerable to membrane cholesterol reduction. **A**, background-subtracted sample images of Filipin staining. pHTm fluorescence was preserved and used to identify transfected cells. AM1-43 was used to identify neurites and for normalization. **B**, quantification of Filipin staining. **B1**, background subtracted filipin signal, One-way ANOVA was used to compare the three conditions and significant variance was detected (F (2, 911) = 28.349, *p* < 0.0001). Cells transfected with I703A have a significantly lower filipin signal compared to WT according to Dunnett’s multiple comparisons test (****, *p* = 0.0001), but there was no difference between G700A and WT (ns, *p* = 0.9908). **B2**, mean normalized Filipin signal (Filipin/AM1-43) in transfected process segments. Dunnett’s multiple comparisons test, performed after one-way ANOVA (F (2, 911) = 89.127, *p* < 0.0001), demonstrates that both G700A (**, *p* = 0.004) and I703A (****, *p* = 0.0001) are different from WT. n_WT_ = 264, n_G700A_ = 206, and n_I703A_ = 444, where n is the number of neurite segments from 3 FOVs of each groups. **C**, sample pHTm images of transfected cells with or without MβCD treatment. Fields of view were not excluded based on failure to respond to NH_4_Cl (provided that they responded to pH 5.5), because dead or otherwise compromised cells and synapses are intended to be included in this analysis. Scale bar, 10 μm. **D**, average surface vs. total pHluorin ratios of WT and mutants with or without MβCD treatment. Two-way ANOVA detected a significant difference after MβCD (F (1, 1131) = 7.542, *p* = 0.0061). There was interaction (F (2, 1131) = 9.205, *p* = 0.0001) but no effect due to the mutation itself (F (2, 1131) = 0.3774, *p* = 0.6858) detected in this data set. Sidak’s multiple comparisons test showed that MβCD significantly increased the ratio in WT and I703A only (WT, *, *p* = 0.0421; G700A, ns, *p* = 0.2037; I703A, ***, *p* = 0.0002). E, average surface vs. total pHTm ratios of two genotypes with or without MβCD treatment. Twoway ANOVA detected significant differences between treatments (F (1, 1131) = 189.1, *p* < 0.0001) and genotype (F (2, 1131) = 38, *p* < 0.0001), and there was an interaction (F (2, 1131) = 6.125, *p* = 0.0023). Sidak’s multiple comparisons test showed that MβCD had a significant effect on the ratio for all genotypes (WT, ****, *p* < 0.0001; G700A, ****, *p* < 0.0001; I703A, ****, *p* < 0.0001). **F**, mean area of synapses. Two-way ANOVA found a significant interaction between the mutation and treatment (F (2, 1131) = 12.79, *p* < 0.0001) and singificant variance based on genotype (F (2, 1131) = 10.27, *p* < 0.0001) and on treatment (F (1, 1131) = 10.38, *p* = 0.0013). Sidak’s mutliple comparisons test showed, in fact, that the MβCD’s effect on synapse size is specific to the mutants (G700A, ****, *p* < 0.0001; I703A, **, *p* = 0.0039) and does not occur in the WT (ns, *p* = 0.1163). **G**, mean roundness of synapses. Two-way ANOVA found a significant interaction between the mutation and treatment (F (2, 1131) = 6.12, *p* = 0.0023), but no singificant difference between the WT and mutants (F (2, 1131) = 0.8548, *p* = 0.4256) or with treatment (F (1, 1131) = 0.4782, *p* = 0.4894). Sidak’s mutliple comparisons test showed, in fact, that the MβCD’s effect on roundness is specific to G700A (**, *p* = 0.0029) and does not occur in the WT (ns, *p* = 0.896) or I703A (ns, *p* = 0.48). For **D-G**, n_WT_ = 131, n_WT+MβCD_ = 135, n_G700A_ = 182, n_G700A+MβCD_ = 235, n_I703A_ = 198, n_I703A+MβCD_ = 256, where n is the number of ROIs corresponding to SypHTm-marked synaptic boutons. **H-J**, average fluorescence changes during 90-min 1 mM MβCD treatment. n_WT_ = 48 ROIs; n_G700A_ = 59 ROIs; n_I703A_ = 53 ROIs.. Shadows represent s.e.m.

Since cholesterol is critical for neuronal membrane integrity ^57, 58, 60^ and functionally essential for the origination and recycling of synaptic vesicles ^61, 62^, we transfected cultures with WT and mutation-bearing SypHTm:T2A:pH-APP-BFP2 and used 100x objectives to visualize morphological details at distal neurites and synaptic boutons (**Figure 6C**). While the transfection efficiency did not differ, there were fewer transfected neurons that survived after the MβCD treatment in both mutant groups than in the WT group (not shown). Using live-cell imaging and ratiometric analysis (i.e. pHluorin out *vs*. total), we found that I703A, like WT, remained responsive to MβCD but G700A did not (**Figure 6D**). In comparison, the pHTm out vs. total ratio exhibited a significant increase for both mutants and WT (**Figure 6E**), matching the previous observation. Importantly, for both mutants but not for WT we observed severe synaptic deterioration in the pHTm channel, including swelling, detachment and failure to respond to NH_4_Cl in both mutants after MβCD treatment (**Figure 6C**). Synaptic swelling is much worse in G700A than I703A-expressing neurons after MβCD treatment, with a ∼59% and ∼38% increase respectively (**Figure 6F**). Furthermore, only G700A demonstrated a significant difference in terms of shape after treatment (**Figure 6G**). Overall, the expression of either mutation caused a severe response to cholesterol depletion likely detrimental to the function and survival of the neurons.

To understand how the synaptic deterioration occurred during the MβCD treatment, we performed time-lapse imaging and analyzed the fluorescence intensities of pHluorin, BFP2 and pHTm (**Figure 6H-J**). Notably, both pHluorin and pHTm signals represented surface as well as partially quenched intracellular proteins in the normal Tyrode’s solution. During the course of 90-minute MβCD treatment, synaptic boutons in WT exhibited a mild decrease of pHTm and BFP2 signal and a moderate increase of pHluorin, which agreed with previous results (**Figure 4**). In the case of G700A, we observed a continuous increase of pHTm and pHluorin signals during the first half of the treatment and it was halted during the second half. The BFP2 signal exhibited considerable fluctuation and a very small overall increase. In conjunction with the observed changes in synaptic morphology, we conclude that the increase of pHTm and pHluorin signals likely represent synaptic swelling and the subsequent cessation of this effect likely reflects synaptic breakdown. In the case of I703A, there was almost no change in pHTm and BFP2 but a larger and longer increase of pHluorin until the end of the treatment, which was in good agreement with the observed morphological changes, indicating synaptic swelling but not breakdown. The time-lapse results match the morphological data, suggesting that G700A caused more severe dominant negative effect than I703A did. The membrane vulnerability caused by G700A could allow more penetration of Filipin, a membrane disrupter ^63^ similar to MβCD, and thus more staining than that of I703A, which was mitigated by membrane density normalization (**Figure 6B**). Potentially, G700A could mask its own change upon the moderate MβCD treatment because the plasma membrane was already disrupted (**Figure 6D&J**). Given the relatively smaller changes in pHTm signals, we further postulate that mutant-associated synaptic swelling was more likely due to surface membrane expansion or the surface deposition of endosomes and/or lysosomes.

## Discussion

In our study, multi-channel fluorescence imaging and ratiometric analysis reveal a hidden link between APP and cholesterol by enabling a quantitative study of APP distribution in the membranes of intact synaptic boutons of live neurons. These studies took advantage of the unique properties of the triple-fluorescence reporter, SypHTm:T2A:pH-APP-BFP2. Despite two fluorescent proteins fused to its N- and C-termini, pH-APP-BFP2 was distributed and cleaved like endogenous APP. The co-expression of SypHTm not only provided us with a landmark for synaptic boutons but also allowed us to monitor neuronal activity and to evaluate the potential interplay between APP and SVs.

While it is well documented that APP is abundant at presynaptic terminals ^18^, the possibility that its surface turnover and cleavage are coupled to SV turnover remains controversial ^17, 64, 65^. Our data showed that APP externalization and subsequent internalization at synaptic boutons was significantly delayed and not quantitatively correlated to SypHTm turnover, suggesting that APP may be localized to the presynaptic active zone and endosome/lysosome membranes instead of specific to SV membranes. The final near-complete retrieval of externalized APP along with minimal changes in surface levels of APP under αS inhibition do suggest that the APP level at the synaptic surface is tightly regulated. However, neuronal activity does not seem to be a major regulator for synaptic APP distribution given the lack of correlation between stimulations and the lateral movement of BFP2 puncta, between APP turnover and SV recycling, and between surface APP and prolonged network activity changes. In light of the recent imaging studies of somatodendritic APP and BACE-1 ^30^, we speculate that the activity-dependent proteolysis of APP/CTF mostly occurred at somatodendritic regions whereas APP/CTF at the distal synaptic boutons behaves and functions differently. Combining pH-APP-BFP2 with red fluorescent protein-tagged secretases like BACE-1 will be useful in addressing this difference.

Cholesterol has been suspected to be a pathological factor in AD, although it remains unclear whether it is cholesterol in circulation, in cell membranes, or in both that matters ^66^. The fact that our moderate MβCD treatment significantly reduced the total pHluorin fluorescence is in good agreement with the notion that cholesterol regulates Aβ production by modulating secretase cleavage ^67^. However, MβCD or membrane cholesterol seems to have a direct effect on surface APP distribution, because cholesterol-dependent change of secretase activities cannot explain the increase of surface APP fraction after MβCD treatment, and because point mutations within the APP cholesterol-binding motif significantly affected its surface distribution independent of γ-cleavage. In addition, cholesterol-induced changes in SV exo-/endocytosis ^61, 68, 69^ could not be the major reason either because the increase of surface APP did not match that of surface SypHTm proportionally. Instead of those indirect mechanisms, we think that MβCD or membrane cholesterol has a more prominent and direct impact on surface-internal trafficking of APP via a recently identified cholesterol-binding motif ^46^. Our tests using two different point mutations have demonstrated that APP’s cholesterol affinity is very important for maintaining the surface APP fraction, either facilitating APP internalization or preventing its externalization independent of SV turnover. γS inhibition did little to increase the surface fraction of mutant APP, further demonstrating that cholesterol directly regulates APP surface distribution. More intriguingly, both mutants rendered the mild MβCD treatment more harmful to neurons despite the presence of endogenous WT APP (i.e. in a dominant negative fashion). The explanation for this phenomenon may reside in APP’s ability to homo- or heterodimerize with APP or CTF respectively. In fact, the cholesterol-binding motif partially overlaps with the proposed dimerization motif ^46, 70^, and competition between cholesterol-binding and APP/CTF-dimerization has been observed in a biophysical study ^59^. Hence, it is possible that the two mutants dimerize with endogenous APP or CTF to render them insensitive to membrane cholesterol change. Additionally, APP-CTF heterodimerization may also be accountable for the increase of surface APP after γS inhibition ^71^.

Our results support the idea that APP is essential for maintaining synaptic membrane cholesterol homeostasis and for the integrity of synaptic membranes, which can be helpful to reconcile many different functions proposed for APP because membrane cholesterol has a very broad impact on various cellular processes including most transmembrane signaling and membrane trafficking pathways ^72, 73^. We postulate that APP is a cholesterol regulator for cholesterol homeostasis between presynaptic surface and intracellular membranes (**Figure 7**). It is well-documented that presynaptic terminals, especially the SVs within, have a much higher concentration of cholesterol in comparison to other parts of neuronal surface membranes ^45^, whereas other intracellular organelles such as endosomes have far less membrane cholesterol. This means that activity-evoked SV release will lead to a substantial increase of surface cholesterol and a correspondingly profound decrease in intracellular membrane cholesterol at synaptic boutons. Furthermore, since synaptic boutons are often far away from the neuronal soma, a local regulatory mechanism of membrane cholesterol such a mediation by APP may be more important for neurons than other cells. Another intriguing fact is that APP resembles the well-studied sterol regulatory element binding protein (SREBP) ^74^ in several aspects including intracellular membrane localization, cholesterol-sensitivity, cleavage by regulated intramembrane proteolysis, and the function of their proteolytic products (i.e. transcription factor for cholesterol metabolism genes). Based on our observation about APP’s externalization in relationship to neuronal activity, γ-secretase inhibition, MβCD, and its own cholesterol-binding site, we further propose that APP can be a multifunctional player by retrieving surfaced membrane cholesterol, balancing intracellular membrane turnover and sensing endosomal cholesterol (**Figure 7**), all of which agree well with APP’s abundance in both surface and intracellular membranes at synaptic boutons. Additionally, the mutual regulation between APP and lipoprotein receptor (LRP1) ^71, 75^ also aligns well with this model. While our study demonstrates the physiological relevance of APP-cholesterol binding and the functional significance of APP’s cholesterol-binding motif, it also raises many interesting questions about the relationship between membrane cholesterol, APP and secretases. More importantly, our observation of increased synaptic vulnerability and membrane disintegration in mutant-expressing neurons implies that the pathological consequences of APP defects may be mediated by membrane cholesterol and may be aligned with the initial synaptic dysfunction prominent in early stage of AD. Given the increasing capabilities of cellular imaging technology, there is now the opportunity to test these questions in CNS neurons, both *in vitro* and *in vivo.* Indeed, with the help of optogenetics, these questions can and should be explored in the context of synaptic activity and network connectivity, which will not only reveal the intrinsic function of APP but also its etiological relevance to AD.

**Figure 7.**
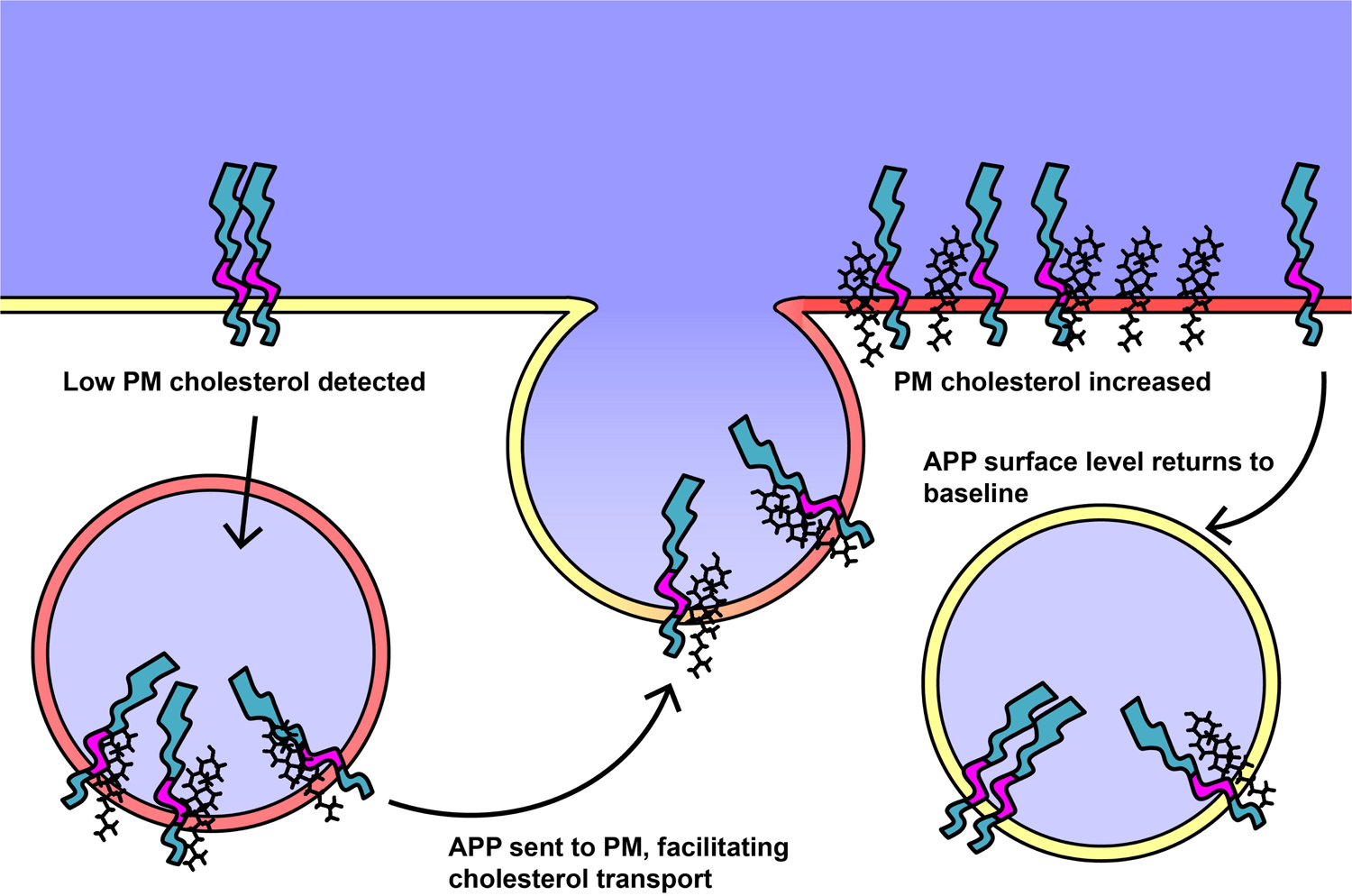
Model for co-modulation of APP and presynaptic membrane cholesterol.

## Acknowledgements

We thank K. E. Kitko for technical help and sharing cell cultures. We thank R.D. Blakley and K.P.M. Currie for comments and discussions. The pHluorin-APP was provided by J. Klingauf lab. We thank all members of the Zhang laboratory for their support. This work is funded by NIH (OD00876101 and NS094738 to Q.Z., and R01 AG056147 to C.R.S.).

## Author Contributions

C.E. DelBove and Q. Zhang conceived this project, designed experiments, analyzed the data and wrote the manuscript. C.E. DelBove conducted all experiments, with two exceptions. R.M. Lazarenko conducted the electrophysiological experiments. C.E. Strothman conducted SypHTm and pH-APP-BFP2 imaging with acute application of 90 K. C.E. DelBove, C.R. Sanders and Q. Zhang conceived and designed the APP mutation experiments. H. Huang generated the APP point mutations. We thank X. Deng for experimental help, and all Zhang laboratory members for discussion.

## Experimental Procedures

### DNA recombination

The pHluorin-APP plasmid, which contains a human synapsin1 promoter, was a gift from Dr. Jürgen Klingauf ^17^. The Synaptophysin-pHTomato plasmid (pTGW-UAS-SypHTm-T2A-BFP2) was a gift from Dr. Yulong Li ^32^. The SypHTm fragment was amplified from the Synaptophysin-pHTomato plasmid and inserted into a mammalian expression vector (pCDNA3.1) containing human synapsin1 promoter using Gibson Assembly Cloning Kit (NEB) ^76^. SypHTm along with the T2A linker was amplified from the pTGW-UAS-SypHTm-2ABFP2 with Phusion kit (NEB) and using 5’- CGTGCCTGAGAGCGCAGTCGAATTAGCTTGGTACCATGGACGTGGTGAATCAGCT GGTGG -3’ (forward primer) and 5’-CCAGGCTGGGCAGCATGGTGGCGGCGGATCCAGGGCCGGGATTCTCCTCCACGT CAC-3’ (reverse primer). SypHTm:T2A:pH-APP was verified by sequencing. Next, BFP2 was amplified from pTGW-UAS-SypHTm-T2A-BFP2 using 5’- CAAGTTCTTTGAGCAGATGCAGAACGCAGCGGCCGCAATGGTGAGCAAGGGCGA GGAGC -3’ (forward primer) and 5’-CATTTAGGTGACACTATAGAATAGGGCCCTCTAGATTACTTGTACAGCTCGTCCAT GCCG -3’ (reverse primer), and was inserted to the C-terminal of pHluorin-APP before the stop codon using the Gibson Assembly Cloning Kit (NEB). The resulting plasmid, SypHTm:T2A:pH-APP-BFP2 was verified by DNA sequencing.

### Cell Culture and Transfection

All animal protocols were approved by the Vanderbilt University Animal Care and Use Committee. Rat postnatal hippocampal cultures were prepared according to an established protocol ^77^ with slight modifications. Briefly, rat hippocampi (CA1-CA3) dissected from P0 or P1 Sprague-Dawley rats were dissociated via an 11-min incubation in Trypsin-EDTA (Life Technologies) followed by gentle trituration using three glass pipettes of different diameters (∼1 mm, 0.5 mm, and 0.2 mm), sequentially. Dissociated cells in suspension were recovered by centrifugation (x 200 g, 5 minutes) at 4°C and re-suspended in plating media consisting of Minimal Essential Medium (MEM, Life Technologies) and (in mM) 27 glucose, 2.4 NaHCO3, 0.00125 transferrin, 2 L-glutamine, 0.0043 insulin and 10%/vol fetal bovine serum (FBS, Omega). 100 μl of resuspended cells were plated onto single round 12mm-ø glass coverslip (∼200,000 cells/mL) pre-coated with Matrigel (Life Technologies) and all coverslips were placed in 24-well plates (ThermoScientific). Cells were allowed to adhere to the coverslips for 30-60 minutes before the addition of 1 mL plating media per well. After 1-2 days in culture, an additional 1 mL media containing (in mM) 27 glucose, 2.4 NaHCO_3_, 0.00125 transferrin, 0.5 L-glutamine, 2 Ara-C, 1 %/vol B27 supplement (Life Technologies) and 5 %/vol FBS was added into every well. Ara-C in the culture media efficiently prevented the overgrowth of astroglia. Calcium phosphate transfection was performed at 8-9 days and most experiments were performed at 14-17 days after the full maturation of neuronal synapses.

### Immunocytochemistry

After treatments or imaging, coverslips were fixed in PBS containing 4% paraformaldehyde for 20 minutes, permeabilized with 0.25% Triton X-100 for 10 minutes and blocked for at least one hour with goat serum and BSA solution all at room temperature (substituting horse serum if a goat primary was used). Next, they were incubated with diluted primary antibodies (see Supplementary table S1) overnight at 4 °C or at room temperature for at least one hour. After incubation with primary antibodies, cells were thoroughly washed and then incubated with specific secondary antibodies labeled by distinct fluorophores (see **Supplementary Table S1**, 1: 1000 dilution for all, Life Technologies or Biotium) at room temperature for at least one hour before mounting.

### Filipin and AM1-43 dual staining

Coverslips were removed from their incubator and washed briefly in pre-warmed (37°C) 4K Tyrode. AM1-43 was diluted in 90K solution and applied to the cells for 1 minute at room temperature. After at least 2 gentle washes with room temperature PBS, the cells were fixed for 30 minutes in 4% paraformaldehyde. The paraformaldehyde was quenched with 1.5 mg/mL glycine in PBS for 10 minutes at room temperature before washing in PBS. Filipin solution (0.05 mg/mL in PBS) was applied to the cells. In order to minimize Filipin permeabilization of the cells, the staining was performed at 4°C for 30 minutes only. The coverslips were then washed and mounted.

### ELISA

The Mouse/Rat sAPPα (highly sensitive) Assay Kit (IBL #27419) was used to measure the sAPPα concentration in the undiluted media. Serum-containing culture media was removed and replaced with 1mL of serum-free NeuroBasal media supplemented with 2 mM L-glutamine, 2% B27-supplement and 100 ng/mL BDNF. Treatments were applied during the media exchange. After 24-hour incubation, the media was collected. Aβ40 was measured using SensoLyte^®^ β-Amyloid (1-40) ELISA kits (AnaSpec). To achieve greater sensitivity for Aβ40 measurement, only 0.5 mL of media was added at time of treatment. Once collected, a protease inhibitor cocktail (Roche) was added to the samples to protect them while they were concentrated to 300 μL using an evaporator. All samples were aliquoted and frozen at -80°C if not used fresh and were only thawed once. After following the instructions provided with each kit, the plates were scanned using a GloMax Discover (Promega). A sigmoidal dose-response curve (constrained to bottom = 0 because of subtraction of media-only blanks) was fit to the standards using Graphpad Prism 7.03 for Windows (Graphpad Software, La Jolla California USA, www.graphpad.com) and the curve was used to calculate the concentration of the samples. 2 technical replicates (2 wells) per biological replicate (media sample) were normalized to the vehicle (DMSO) control and averaged together before the three biological replicates were averaged for an n of 3. Samples slightly below background with a technical replicate slightly above background were treated as 0 rather than excluded. The experiments in **Supplementary Figure 6** were performed slightly differently. BCC or NBQX treatments were added to the existing media 2 hours before collection, so that the result measures the effects of both Aβ and sAPPα secretion and degradation in a physiological environment. An additional sample was taken immediately following application of the vehicle control (H_2_O), which we termed the 0 hour sample. We discovered that NBQX treatment interfered with the Aβ40 ELISA and reduced the signal. In order to correct for this, we fit a second order polynomial curve to samples with and without NBQX added immediately before adding them to the wells. This correction was then applied to all NBQX-treated samples using Prism. After the correction, all samples were normalized to 0 hour.

### Immunofluorescence imaging and analysis

All ICC imaging was performed on a Nikon Eclipse Ti inverted microscope equipped with a 100X Plan Apo VC objective (N.A. 1.40) and an Andor iXon+ 887 EMCCD camera. All imaging settings including power of the excitation light source, fluorescence filter sets (excitation, dichroic and emission filters, all from Semrock), exposure time and EM gain were kept consistent for imaging different groups with the same immunofluorescence labeling. The optical filter sets (Chroma and Semrock) for Alexa 405/DAPI, 488, 568, and 647 fluorescence were, respectively: Ex 405/20X, DiC 425LP and Em 460/50; Ex 460/50, DiC 495LP and Em 535/25; Ex 565/25, DiC 585LP and Em 644/90; Ex 644/10 DiC 660LP and Em 710/50. Image analysis was performed in Fiji ^78^, a distribution of ImageJ ^79^. First, four cell-free areas were selected as background regions and their mean fluorescence intensities were averaged to obtain an overall background fluorescence intensity value. Process ROIs were hand drawn in areas with a relatively low background. Synapse ROIs were hand-drawn from all synaptophysin1 puncta along the selected processes. Only synapse ROIs with an APP signal above background were quantified in most cases. For the purposes of defining nonsynaptic regions for quantification, all areas with a high Syp signal were subtracted from the process ROIs and the remainders were divided into sections of similar area. To avoid bias towards any field of view, an equal number of ROIs from each FOV (equal to the least number of ROIs per FOV) was randomly selected and pooled. The n for each condition is the total number of pooled ROIs. Background subtraction and random selection of ROIs was performed in Microsoft Excel. The values were imported into Prism 7.03 for plots and statistics. Statistical tests were chosen based on the experiment.

Object-based colocalization (**Supplementary Figure 1**) was performed using the Synapse Counter plug-in for ImageJ ^80^. No alterations were made to the images except to exclude transfected somas. Briefly, for 512x512 16bit images taken at 100x, the rolling ball radius was set to 8, the maximum filter radius to 1.0, Otsu threshold adjustment was used and minimum and maximum particle sizes were set to 5 and 400. For the purpose of this quantification, fields of view were used as the n rather than regions of interest as in other quantifications.

Filipin imaging (**Supplementary Figure 7**) was performed using AM1-43 to select FOV to minimize photobleaching in the Filipin channel. Due to the lack of synaptic markers, ROI selection was focused on the processes. Rather than hand drawing all processes as in quantifications that feature transfected cells only, the images were thresholded as a base, additional processes of lower intensity were added by hand, and the somas and debris were excluded. The processes were divided into ROIs of relatively uniform size. Areas with a lower density of astrocytes were intentionally selected but astrocytes could not be completely separated from the neurons.

### Live cell imaging and analysis

Live cell imaging was performed with a Nikon Eclipse Ti inverted microscope, a 20X or 100X Plan Apo VC objective and an Andor iXon+ 887 EMCCD camera or a Hamamatsu Flash4 (**Figure 1D**). 12 mm coverslips were mounted in an RC-26G imaging chamber (Warner Instruments) bottom-sealed with a 24x40 mm size 0 cover glass (Fisher Scientific). The chamber was fixed to a PH-1 platform (Warner Instruments) placed on the microscope stage. Gravity perfusion-powered solution exchange was controlled by a VC-6 valve control system and a 6-channel manifold (Warner Instruments) with a constant rate of ∼50 μL/sec which allowed a complete bath solution turnover in the recording chamber in under 30 s. Image acquisition and synchronized perfusion were controlled via Micro-manager software. For every fluorophore, the acquisition settings including excitation power, fluorescence filter set (excitation, dichroic and emission filters), exposure time, camera gain and frame rate were all kept the same among different samples on all experiments. The optical filter sets (Chroma and Semrock) for Alexa 405/BFP2, pHluorin and pHTomato were, respectively: Ex 405/20X, DiC 425LP and Em 460/50; Ex 480/20X, DiC 495LP and Em 535/40; Ex 560/40M, DiC 585LP and Em 610/20nm BP. Samples were exposed to normal Tyrode’s saline at pH 7.35, a 50mM NH4Cl solution and normal Tyrode’s solution adjusted to pH5.5 sequentially. Some of the older data sets in **Figure 4** and **Figure 2** were collected using NH_4_Cl before 4K. Tyrode’s saline contains (in mM): 150 NaCl, 4 KCl, 2 MgCl_2_, 2 CaCl_2_, 10 N-2 hydroxyethyl piperazine-n-2 ethanesulphonic acid (HEPES), 10 glucose, pH 7.35 or pH 5.5. The 50 mM NH_4_Cl solutions was made by substituting for NaCl equimolarly, pH 7.35. All solutions contained 10 μM NBQX and 20 μM D-AP5 except in the experiment in Figure 7 F-H, which was meant to replicate the experiment in **Figure 7 A-E** during which MβCD was applied to the home media as a pretreatment in the absence of such activity altering drugs. StackReg ^81^ was used to correct for state drift. MultiStackRegistration (Brad Busse) was used to align the pHluorin stacks using the pHTm stacks as reference, with individual frames adjusted as needed manually. For the purposes of quantification, the 3 most representative frames (5 for **Figure 4** and **Figure 2**) were chosen in each solution and averaged together in ImageJ. For example, as NH_4_Cl is applied, fluorescence increases to a peak and then begins to decrease. Three consecutive frames with the highest fluorescence signal are used to represent of the maximum fluorescence. Because BFP2 signal is not pH-dependent, all BFP2 frames were averaged together for quantification. Using averaged frames rather than taking the beginning, maximum and minimum from each trace reduces the error caused by actively moving puncta passing by a synapse. Average frames were used for all analyses except **Figure 3**, which is intended to detail acute responses to activity rather than to determine the surface and internal fractions of the fluorophores at a steady state as in the other figures. For live cell imaging analysis, synapses were defined based on the SypHTm signal; specifically, all static puncta. ROIs for synapses and neurites were manually selected in ImageJ. In addition, multiple cell-free areas were selected as background ROIs. Separate background ROIs were selected for BFP2 because astrocytes and untransfected neurons produced a high background in that channel; some ROIs chosen based on SypHTm were excluded due to the background in the blue channel in order to avoid individually defining the background for each ROI.

Nonsynaptic ROIs were defined by manually drawing processes as seen on all 3 channels, subtracting out any regions suspected to be synapses, and dividing the remainder into similarly sized ROIs. Additionally, only ROIs with a detectable SypHTm signal in normal Tyrode’s solution were included because puncta that are only visible in NH_4_Cl are not necessarily synapses. For the purposes of defining nonsynaptic regions for quantification, all stationary SypHTm puncta were subtracted out. The somatodendritic regions were excluded, usually by choosing an imaging field without a transfected soma, due to high background and low prevalence of probable synapses compared to puncta visible only in NH_4_Cl. The values of all ROIs, including background, for each image (average NH_4_Cl, 4K, pH5.5 for SypHTm and pHluorin and stack average for BFP2) were exported to Microsoft Excel. The background was then subtracted, and then NH_4_Cl – 4K was used to calculate the internal protein and 4K – 5.5 was used to calculate the surface protein. For the purposes of quantification of APP in the synapses, only ROIs with a positive value for Total, Internal and Surface in the pHluorin and pHTm channels were included in the analysis. If the BFP2 signal was not higher than background, the ROI was also excluded. To avoid bias towards any field of view, an equal number of ROIs from each FOV was randomly selected and pooled. The n for each condition is the total number of pooled ROIs.

Kymographs were generated by manually tracing the processes using the Segmented Line tool in Fiji based on an averaged image of the BFP2 channel, using the "Reslice" command (Output Spacing = 1 pixel, Avoid interpolation). Paths of the moving puncta were traced by hand using the Segmented Line tool and saved to the ROI Manager.

These selections were exported as XY coordinates with interpolation of 1 pixel. In order to be as accurate as possible, the actual time each image was taken was extracted and used to convert vertical pixel distances to time because the timing did not sync perfectly to the frame number across different fields of view. Because puncta sometimes split, appeared and disappeared, the n was determined separately at each time point for the purpose of calculating the mean and SEM. Puncta that disappeared were not counted as a zero but as a blank, while puncta that were still present but were not moving were included in the analysis. Synapses were determined using SypHTm puncta that did not move across the smaller number of frames collected of the channel. For each kymograph, the position of all synapses along the processes was listed on each spreadsheet page. Using Microsoft Excel’s array functions, the closest synapse to the puncta at any given time point was selected from the list. When a punctum moving between two synapses A and B became closer to B than A, it was then considered to be moving in the positive direction towards the nearest synapse. The position of the puncta with respect to the soma was not considered for the purpose of this calculation because the experimental question was whether lateral movement toward or away from the synapse. There was no marker used in any live cell imaging that can separate axons from dendrites, so no claim is made about the identity of the processes quantified. However, due to the distance from the soma needed to get lower backgrounds, the narrow and consistent width of the selected processes, and the expression pattern of SypHTm, they are most likely axons.

The analysis of **Figure 7A-D** was performed differently because the size was the most important feature to be quantified, and hand drawn ROIs were therefore unsuitable. The images were also collected differently (detailed below). First, Convoluted Background Subtraction from the BioVoxxel Toolbox (http://imagej.net/BioVoxxel_Toolbox) was used, followed by Non-local Means Denoising ^82, 83^, bandpass filter, auto-local threshold using the Bernsen method, Adjustable Watershed (Michael Schmid) with a tolerance of 0.1. Using a macro, the same procedure was applied to all images and then debris were manually removed, non-static ROIs were excluded, and compounded ROIs that escaped the watershed were manually separated. The measure tool was then used to collect the data. All of the intensity values were taken from the original, unaltered images. During the collection of this data, multiple fields of view were observed from the same coverslip between solution changes using the Position List feature on MicroManager to return to the locations. 10 frames were collected from each channel at each field of view in each solution, aligned and averaged. To correct for cumulative photobleaching, values for NH4Cl and pH 5.5 were multiplied by arbitrary factors so that the quantification can still be performed. Furthermore, due to morphology changes over time during this extended imaging, the ROIs were manually moved as the processes moved based on the pHTm channel, which the other channels were aligned to. Additionally, due to the high signal to noise ratio each ROI had its own background ROI defined by shifting all ROIs in the manager about 10 pixels over in the x and y directions and manually moving the background ROIs from there to the most suitable area with a similar background (for example, a transfected cell-free area on the same auto fluorescing flat glial cell as the synapse). After the background subtraction and arbitrary photobleach correction was equally applied to all conditions, ROIs were excluded as before for having below background signal in any of the channels. Because of this, it is difficult to compare the fluorescence results from this figure to other experiments in the paper directly. For this experiment, it was very important to select FOV without consideration to factors such as the size of the synapses, process health and morphology. Because fields of view were not chosen based on the quantity of synapses but rather on the presence of transfected punctated structures at all, ROIs were not randomly selected from each field of view and instead all were included and pooled.

### Statistical tests

Statistical tests were performed using Graphpad Prism 7.03. Each test is listed in the figure legends, but in general two-tailed unpaired *t*-tests were used to compare two conditions, ordinary one-way ANOVA and Sidak’s multiple comparisons test was used to compare results within one variable in experiments with two variables.

## Supplementary Information

The supplementary materials including supplementary figures and legends, supplementary table and supplementary movies.

